# Parallel natural selection in the cold-adapted crop-wild relative *Tripsacum dactyloides* and artificial selection in temperate adapted maize

**DOI:** 10.1101/187575

**Authors:** Lang Yan, Sunil Kumar Kenchanmane Raju, Xianjun Lai, Yang Zhang, Xiuru Dai, Oscar Rodriguez, Samira Mahboub, Rebecca L. Roston, James C. Schnable

## Abstract

Artificial selection has produced varieties of domesticated maize which thrive in temperate climates around the world. However, the direct progenitor of maize, teosinte, is indigenous only to a relatively small range of tropical and sub-tropical latitudes and grows poorly or not at all outside of this region. *Tripsacum*, a sister genus to maize and teosinte, is naturally endemic to the majority of areas in the western hemisphere where maize is cultivated. A full-length reference transcriptome for *Tripsacum dactyloides* generated using long-read isoseq data was used to characterize independent adaptation to temperate climates in this clade. Genes related to phospholipid biosynthesis, a critical component of cold acclimation on other cold adapted plant lineages, were enriched among those genes experiencing more rapid rates of protein sequence evolution in *T. dactyloides*. In contrast with previous studies of parallel selection, we find that there is a significant overlap between the genes which were targets of artificial selection during the adaptation of maize to temperate climates and those which were targets of natural selection in temperate adapted *T. dactyloides*. This overlap between the targets of natural and artificial selection suggests genetic changes in crop-wild relatives associated with adaptation to new environments may be useful guides for identifying genetic targets for breeding efforts aimed at adapting crops to a changing climate.

## Significance Statement

Corn was domesticated in Central America and is very sensitive to cold and freezing temperatures. Eastern gamagrass is a close relative of corn, is native to prairies throughout the United States, east of the rocky mountains, the region we now call the corn belt and can survive the winter. We compared rates of protein sequence evolution across the same genes in seven grass species to identify genes likely to be involved in adapting gamagrass to life in the corn belt. We identified a specific metabolic pathway likely involved in cold and freezing tolerance and also found that many of the same genes were targets of selection when humans started developing new varieties of corn to grow in temperate North America.

## Introduction

The common ancestor of maize and *Tripsacum dactyloides* was adapted to a tropical latitude, yet today domesticated maize and wild *T. dactyloides* both grow in large temperate regions of the globe. The adaptation of tropical maize landraces to temperate environments required changes to flowering time regulation and adaption to new abiotic and biotic stresses [1]. As both a leading model for plant genetics and one of the three crops that provides more than one half of all calories consumed by humans around the world, maize *(Zea mays* ssp. *mays)* and its wild relatives have been the subject of widespread genetic and genomic investigations. The closest relatives of maize are the teosintes, which include the direct wild progenitor of the crop (*Z. mays* ssp. *parviglumis)* as well as a number of other teosinte species within the genus *Zea* (Table 1). Outside the genus *Zea*, the closest relatives of maize are the members of the sister genus *Tripsacum* (Figure 1A-H). Together, these two genera form the subtribe *Tripsacinae* within the tribe *Andropogoneae* [2]. Despite their close genetic relationship, some species of the genus *Tripsacum* are adapted to a much wider range of climates than wild members of the genus *Zea* (Figure 1).

**Table 1.** Taxonomic comparison between genus *Tripsacum* and *Zea*

| *Tripsacum*<sup>1</sup> |  |  | *Zea*<sup>2</sup> |  |  |
| --- | --- | --- | --- | --- | --- |
| Species | Life cycle | Ploidy | Species | Life cycle | Ploidy |
| *T. dactyloides* L. (Gamagrass) | perennial | diploid, tetraploid | *Z. perennis* | perennial | tetraploid |
| *T. andersonii* (Guatemala grass) | perennial | diploid, tetraploid | *Z. diploperennis* | perennial | diploid |
| *T. australe* | perennial | diploid | *Z. nicaraguensis* | annual | diploid |
| *T. maizar* | perennial | diploid | *Z. luxurians* | annual | diploid |
| *T. laxum* | perennial | diploid | *Z. m. huehuetenangensis* | annual | diploid |
| *T. floridanum* | perennial | diploid | *Z. m. mexicana* | annual | diploid |
| *T. cundinamarce* | perennial | diploid | *Z. m. parviglumis* | annual | diploid |
| *T. zopilotense* | perennial | diploid |  |  |  |
| *T. bravum* | perennial | diploid |  |  |  |
| *T. latifolium* | perennial | tetraploid |  |  |  |
| *T. pilosum* | perennial | tetraploid |  |  |  |
| *T. lanceolatum* | perennial | tetraploid |  |  |  |
| *T. peruvianum* | perennial | tetraploid, pentaploid, hexaploid |  |  |  |

<sup>1</sup>References for *Tripsacum* are [5, 51].
<sup>2</sup>References for *Zea* are [52, 53, 3, 54, 55].

**Figure 1.**
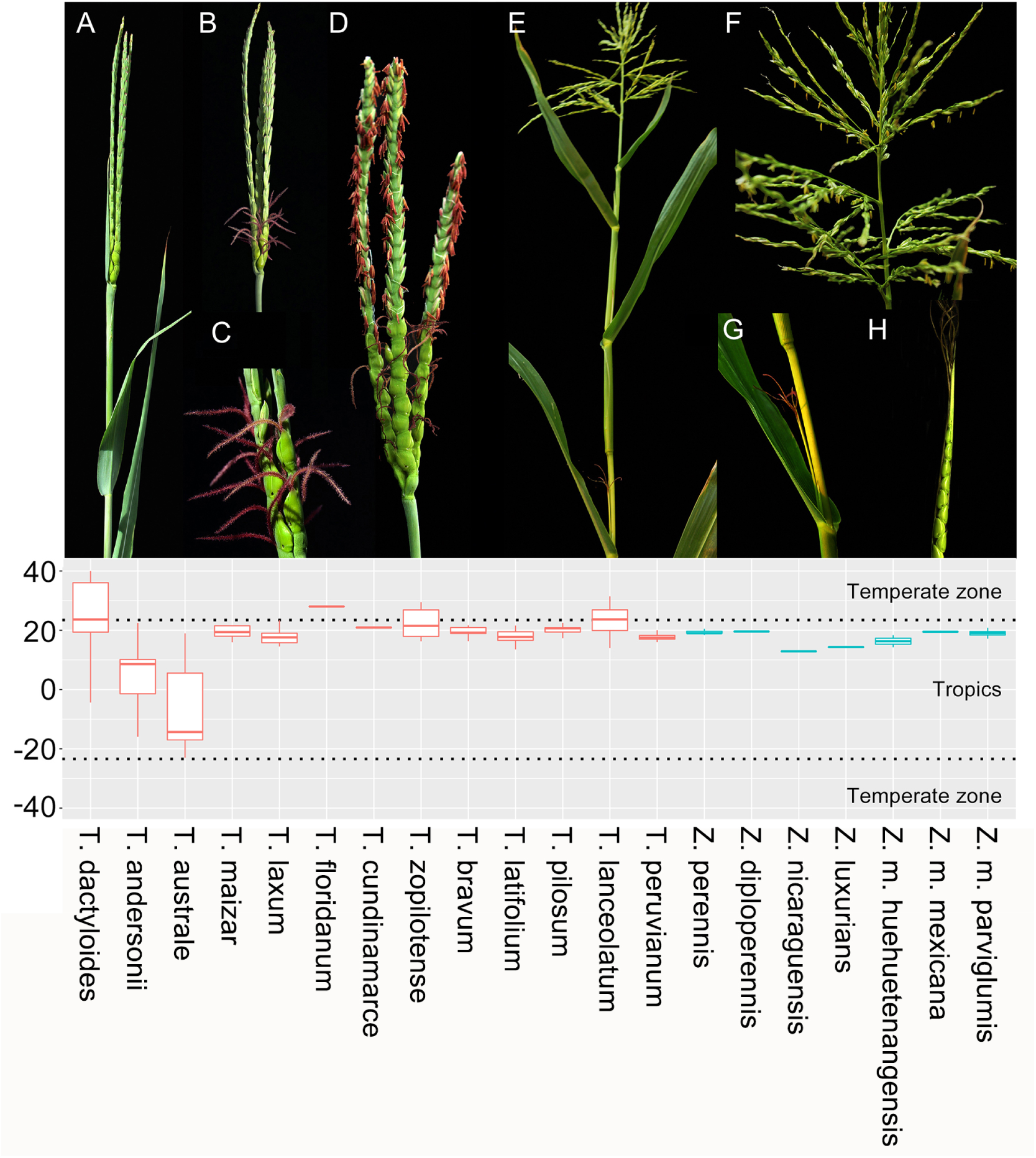
Morphological characteristics of inflorescences for representative species in Tripsacum and Zea. *T. dactyloides* (tripsacum, A-D) and *Zea mays* ssp. *parviglumis* (teosinte, E-H). (A) Early stage inflorescence. (B) Intermediate stage inflorescence with silks exerted from female spikelets. (C) magnified view of female spikelets. (D) mature inflorescence with both silks exerted from female spikelets (base) and anthers exerted from male spikelets (top). (E) Separate male and female inflorescences in teosinte. (F) magnified view of male inflorescence with anthers exerted. (G) magnified view of female inflorescence with silks exerted (H) hard fruitcases surrounding teosinte seeds (absent in domesticated maize). Latitudinal distribution of reported observations in GBIF for wild species within the genus *Tripsacum* (red) and *Zea* (teal). The boundaries of tropical latitudes (ie the tropic of cancer and tropic of capricorn) are marked with dashed black lines. Individual outlier datapoints more than 1.5x the interquartile range are omitted in this display.

The direct progenitor of maize, *Z. m. parviglumis*, is confined to a relatively narrow native range that spans tropical and subtropical areas of Mexico, Guatemala, Nicaragua and Honduras [3]. Other species and subspecies within the genus *Zea* are also largely confined to the same geographic region [4, 3]. In contrast, *T. dactyloides* is widely distributed through temperate regions of both North and South America [5, 6], largely mirroring the distribution of modern agricultural production of maize in the western hemisphere. The common ancestor of *Zea* and *Tripsacum* is predicted to have been adapted to tropical latitudes [5, 7, 8]. Therefore, the study of how natural selection adapted *T. dactyloides* to temperate climates represents an informative parallel to the adaption of maize to temperate climates through artificial selection. *T. dactyloides* is also a potential source of insight into the genetic changes responsible for traits such as disease and insect resistance, drought and frost tolerance, many of which are targets for maize improvement [9, 10, 11].

Until recently, molecular sequence data for *Tripsacum* species was largely been generated to serve as an outgroup for molecular evolution studies in maize (as reviewed [12]). As part of Hapmap2, 8x short read shotgun data generated from *T. dactyloides* [9], additional low pass genomic data has been generated for several other species in the Tripsacum genus [13], and recently, Illumina transcriptome assemblies of two species in the genus Tripsacum - *T. dactyloides* and *T. floridanum* became available [11]. Here we employ PacBio long-read sequencing to generate a set of full length transcript sequences from *T. dactyloides*. Using data from orthologous genes in maize, *T. dactyloides, Sorghum bicolor, Setaria italica*, and *Oropetium thomaeum*, a set of genes with uniquely high rates of nonsynonymous substitution in *T. dactyloides* were identified. We show that a surprisingly large subset of these genes are also identified as targets of selection during artificial selection for maize lines adapted to temperate climates. A specific metabolic pathway identified through this method - phospholipid metabolism - is linked to cold and freezing tolerance in other species and we demonstrate that the metabolic response of this pathway to cold stress shows functional divergence between maize and *T. dactyloides*.

## Results

### Sequencing and analysis of full-length *T. dactyloides* transcripts

PacBio Iso-seq of RNA isolated from a single *Tripsacum dactyloides* plant grown from seed collected from the wild in eastern Nebraska (USA) was used to generate 64,326 HQ consensus sequences (See Supplemental Results; Table S1). These sequences were aligned to the maize reference genome, which identified 24,616 isoforms corresponding to 14,401 annotated maize gene models (See Supplemental Results; Table S2). A total of 1,259 high quality consensus sequences from *T. dactyloides* failed to map to maize reference genome, which was generated from the maize inbred B73. This number is roughly consistent with another report of 1,737 *T. dactyloides* transcripts which lack orthologs in the maize genome [11]. Sequences which lacked apparent homologs in the maize reference genome were aligned to NCBI’s RefSeq plant database. Two-hundred-and-sixty-three of these sequences aligned to genes from other grass species such as *Sorghum bicolor, Setaria italica, Oryza sativa* (Table S3). These genes may either represent genes missed when generating the maize reference genome assembly [14], genes present within the maize pan-genome but absent from the specific line used to generate the maize reference genome [15], or genes present in the common ancestor of maize and *T. dactyloides* but lost from the maize lineage sometime after the *Zea/Tripsacum* split. As a partial test of these different models, these sequences were aligned to the published genome sequence of a second maize inbred (PH207) [16]. Only four of these sequences aligned to the PH207 genome, indicating that the majority of this population of sequences are more likely to represent losses from the maize linage sometime after the *Zea/Tripsacum* split, rather than genes left out of the maize reference genome assembly [14] or genes present within the maize pan-genome but absent from the specific line used to generate the maize reference genome [15].

Patterns of alternative splices observed between multiple *T. dactyloides* transcripts which aligned to the same maize genes followed similar patterns to those identified in studies of alternative splicing in maize based on either short read or long read technology (see supplemental results). However, despite greater sequencing depth and sampling a wider range of tissues, maize Iso-seq dataset did not identify alternative splicing events corresponding to the specific alternative splicing events identified in *T. dactyloides* in 85.7% of cases (2,447 of 2,856 orthologous genes). This result is consistent with the rapid divergence of most AS patterns between even closely related species (see Supplemental Results).

### Identification of *T. dactyloides* genes experiencing rapid protein sequence evolution

A total of 6,950 groups of orthologous genes present in seven grass species were identified using the tripsacum-maize orthologous relationships (see supplemental results), plus an existing dataset of syntenic orthologous genes across six grass species with known phylogenetic relationships (Figure 2A) [17]. These groups included consisted of 4,162 one-to-one, 1,436 one-to-two and 1,352 two-to-two orthologous gene sets due to the WGD event shared by maize and *T. dactyloides*. The overall distribution of synonymous substitution (Ks) values for branches leading to individual species scaled with branch length (Figure 2B). However, while *Zea* and *Tripsacum* are sister taxa, the average maize gene showed more synonymous substitutions than the average *T. dactyloides* gene. In 1,775 cases the branch leading to *T. dactyloides* had the highest Ka/Ks ratio of all branches examined and in 1,114 the branch leading to maize had the highest Ka/Ks ratio (Figure 2C, Figure S2). This bias towards more gene groups showing the highest Ka/Ks values in maize or *T. dactyloides* rather than showing the highest values sorghum, foxtail millet, or oropetium, as well as the presence of more extremely high outlier values in these two species (Figure 2C), likely results from the fact that Ka/Ks ratios are based on a smaller absolute counts of substitutions per gene along shorter branches and therefore exhibit greater variance. In the analyses below, the set of genes experiencing accelerated rates of protein sequence evolution in maize were used as a control set for any analysis of patterns observed in fast evolving *T. dactyloides* genes.

**Figure 2.**
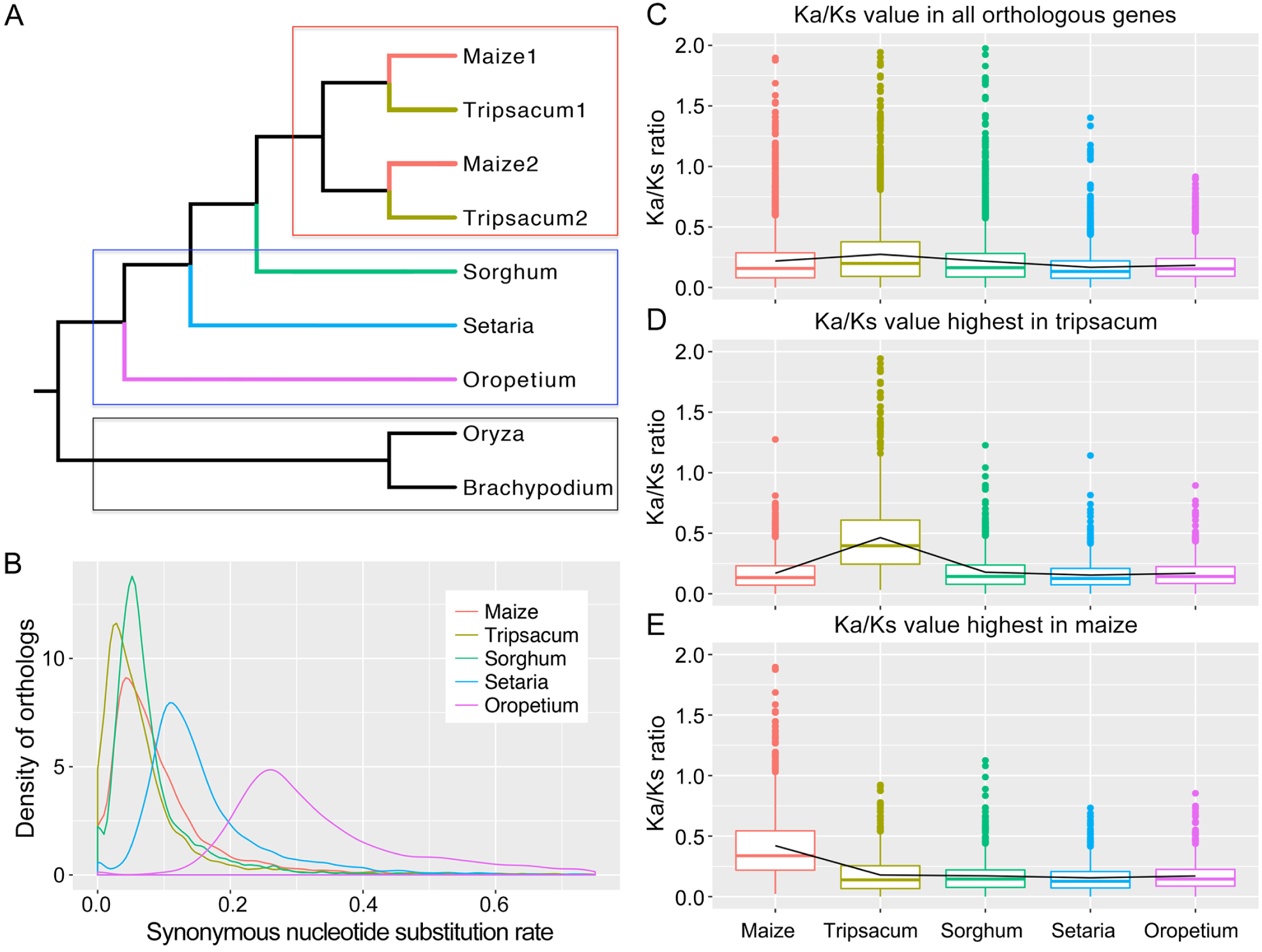
Synonymous substitution rates and ratios of synonymous and nonsynonymous substitution rates across five related grass species.(A) Phylogenetic relationships of the seven species employed in this study. Red, blue and black boxes indicate target species, background species and outgroups respectively. (B) Distribution of synonymous substitution rates (Ks) in orthologous gene groups across each target and background species. Observed synonymous substitution rates in *T. dactyloides* are comparable to or less than those observed in maize. (C) Distribution of Ka/Ks ratios for orthologous genes conserved in maize, *T. dactyloides*, sorghum, foxtail millet (Setaria), and oropetium. (D) Subset of orthologous gene groups displayed in panel C, where the gene copy found in the in *T. dactyloides* transcriptome exhibits a higher Ka/Ks ratio than the same gene in the other four species tested. (E) Subset of orthologous gene groups displayed in panel C, where the gene copy found in the maize genome exhibits a higher Ka/Ks ratio than the same gene in the other four species tested.

Genes with signatures of rapid evolution in *T. dactyloides* tended to be associated with the functional annotations “stress response” and “glycerophospholipid metabolic process”, whereas fast-evolving genes in maize were enriched in the functions microtubule cytoskeleton organization, nutrient reservoir activity and ATPase activity. Figure 3A illustrates the distribution of Ka/Ks ratios in *T. dactyloides* and maize for genes where the branch leading to one of these two species exhibited the highest Ka/Ks ratio among the five species examined. Multiple fast-evolving genes involving in cell response to stimulus and stress had extremely high Ka/Ks ratios (> 1) in *T. dactyloides*, consistent with positive selection for increased abiotic stress tolerance in *T. dactyloides* relative to maize and other related grasses. The annotated functions of the maize orthologs of *T. dactyloides* genes experiencing accelerated protein sequence evolution include cold-induced protein, drought-responsive family protein, salt tolerance family protein, etc. (Table S4).

**Figure 3.**
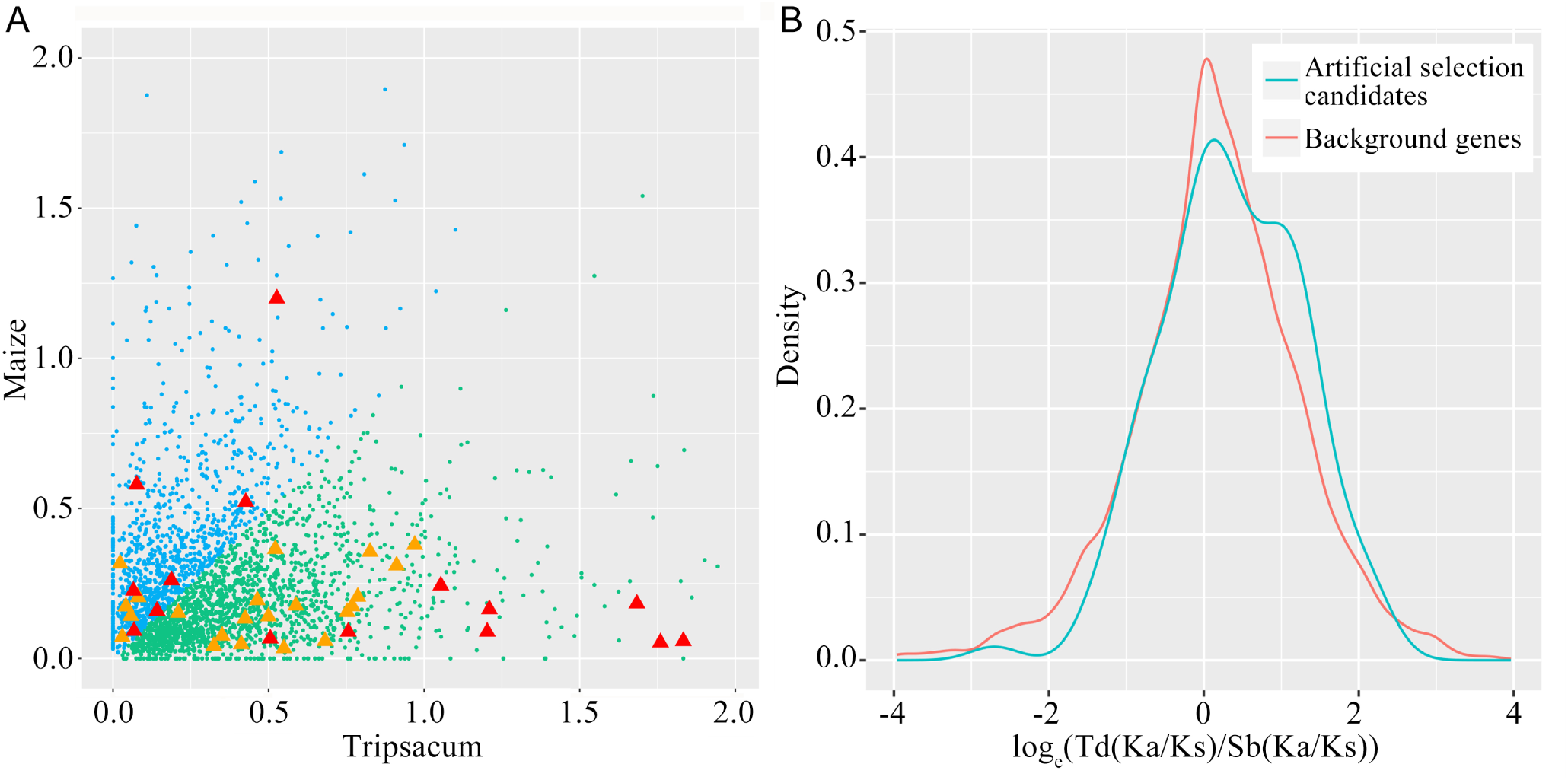
Comparative distribution of Ka/Ks ratios between maize and *T. dactyloides*. (A) The relationship between Ka/Ks ratios observed in maize (blue) and those observed in *T. dactyloides* (green) for genes having the higher Ka/Ks ratio in one of these two taxa than in any of the three background taxa (sorghum, setaria, and oropetium). Orange triangles mark genes annotated as involved in glycerophospholipid metabolism, while red triangles mark genes involved in stress response. (B) Plot of the log transformed ratio of ratios between *T. dactyloides* and sorghum Ka/Ks values for two populations of genes. The artificial selection candidate set includes all orthologous gene groups conserved across the seven species employed in this analysis but where the maize gene copy was identified as a target of artificial selection between tropical and temperate maize inbreds. The background gene set includes all other orthologous gene groups conserved across these seven species. The artificial selection candidate gene set exhibits a skew towards higher ratios of Ka/Ks ratios relative to the background gene set.

### The phospholipid metabolic pathway is a specific focus of accelerated protein sequence evolution in *T. dactyloides*

In the process of identifying *T. dactyloides* genes that might experience accelerated evolution for temperate climate adaptation, we noticed multiple genes annotated as participating in the phospholipid metabolism pathway where the highest Ka/Ks ratio for that gene were observed in the branch leading to *T. dactyloides*. While several genes in this pathway also showed signs of accelerated evolution in maize, the bias towards high rates of protein sequence change in *T. dactyloides* was dramatic (Figure 3A). Using log-transformed Ka/Ks values, genes in the phospholipid biosynthesis pathway exhibited a significantly higher range of Ka/Ks values than the background set of other genes (p-value = 1.87e-04). In contrast, maize genes in the same exhibited significantly lower Ka/Ks values than background maize genes (p-value = 2.38e-04). Comparing the ratio of Ka/Ks values for between same genes in both maize and *T. dactyloides*, genes in the phospholipid biosynthesis pathway showed significantly higher ratios of Ka/Ks than background genes (p-value = 4.16e-05) (Figure S3A).

Phospholipids are a class of lipids that are a major component of cell membranes and include lipids with head groups such as phosphatidate (PA), phosphatidylethanolamine (PE), phosphatidylcholine (PC), phosphatidylglycerol(PG) and phosphatidylserine (PS) which share overlapping biosynthesis pathways and are often inter-convertible (Figure 4a). The set of genes involved in phospholipid metabolism which experienced accelerated evolution in *T. dactyloides* were particularly concentrated in the pathway leading to PC which is the major phospholipid component of non-plastid membranes. PC also tends to control membrane desaturation through acyl-editing [18]. Testing confirmed that *T. dactyloides* seedlings grown from seed collected as part of the same expedition were able to tolerate prolonged 4 °C cold stress while the same temperature stress treatment produced significant levels of cell death in maize seedlings from the reference genotype B73 (Figure S3B-F).

**Figure 4.**
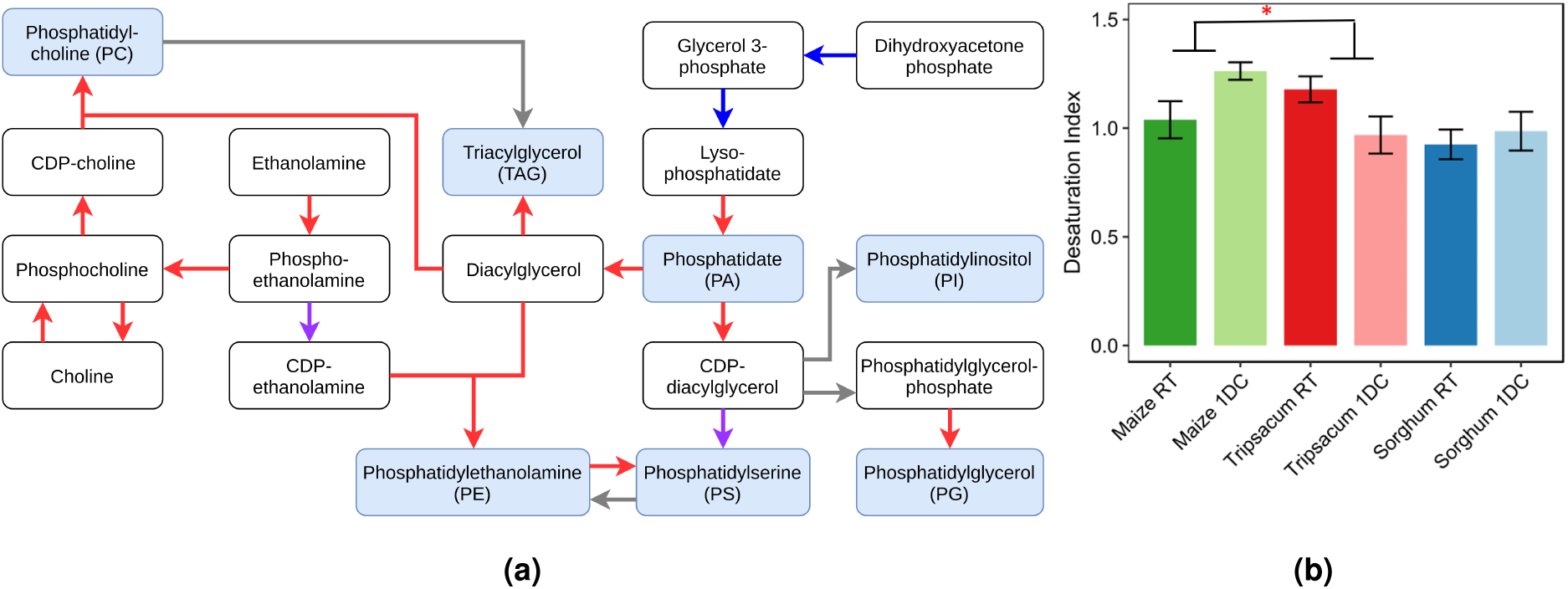
A partial diagram of glycerophospholipid metabolism including phospholipid synthesis. a) Each name enclosed in a white box corresponds to an intermediate product in glycerophospholipid metabolism. Each name enclosed in a blue box indicates a mature lipid quantified in maize and *T. dactyloides*. If at least one gene encoding an enzyme which catalyzes a specific reaction was found to be evolving faster in *T. dactyloides* than any of the other grass species tested, the arrow indicating that reaction is drawn in red. If at least one gene encoding an enzyme which catalyzes a specific reaction was found to be evolving faster in maize than any of the other grass species tested, the arrow indicating that reaction is drawn in blue. If multiple genes encode different isoforms of the same enzyme and at least one was identified in the maize fast evolving gene set and a separate gene was identified in the Tripsacum fast evolving gene set, the arrow is drawn in purple. In cases where none of the genes encoding an enzyme catalyzing a particular reaction were identified as fast evolving in either species, the arrow is drawn in gray. b) Estimation of phosphatidyl-choline (PC) desaturation in maize, sorghum, and *T. dactyloides* before and after one-day of cold treatment. denotes t-test p-value 0.035.

### *T. dactyloides-specific* changes in the response of lipid metabolism to cold stress

RNA-seq and membrane lipid profiles were collected from maize, sorghum, and *T. dactyloides* seedlings under control and cold stressed conditions. The inclusion of sorghum provided a method to ascertain whether maize or *T. dactyloides* represented the ancestral state when metabolic or transcriptional patterns of responses to cold differed between the two species. As previously reported, no GO terms were significantly enriched among differentially regulated orthologs (DROs) of maize and sorghum, three hours after the onset of cold stress. At the same time point, DROs between *T. dactyloides* and maize + sorghum were enriched in genes related to photosynthesis light harvesting, protein-chromophore linkage, and chlorophyll metabolic genes. At 6 hr post-stress, genes related to chloroplast and mitochondrial metabolic processes were differentially regulated in *T. dactyloides* compared to maize and sorghum (Table S5). Specifically, chloroplast and mitochondrial RNA processing, modification and metabolic process genes were up-regulated in *T. dactyloides* at 6 hr post stress, while a suite of abiotic stress responsive genes, annotated as responding to osmotic stress, heat stress or salt stress, as well as genes involved in histone modification and methylation, chromatin organization and ethylene signaling were down-regulated (Table S6). These observations are all consistent with maize and sorghum both experiencing severe impairment of fundamental biological processes within six hours of the onset of cold stress, while *T. dactyloides* seedlings remained relatively healthy under equivalent levels of stress treatment.

The overall pattern changes in the abundances of the major membrane lipids described in (Figure S4A) between control and cold stress conditions were not significantly different across the three species although individual statistically significant changes in lipid abundance in response to cold were observed in each species. However, this assay also allowed the quantification of fatty acid desaturation for individual lipid types. Fatty acids are initially synthesized in a more saturated state. Hence a decrease in desaturation can be an indicator of increases in lipid synthesis. Increase in desaturation can serve as a signaling mechanism and also increases membrane fluidity allowing plants to avoid membrane damage at low temperatures (As Reviewed [19]). Statistically significant changes in lipid desaturation were observed for several lipid classes in individual species (Table S7). However, PC was unique in that the pattern of desaturation change in response to cold was consistent between maize and sorghum, and opposite in *T. dactyloides* (Figure 4b). Fast evolving genes in *T. dactyloides* lipid biosynthesis genes were concentrated in pathways leading to PC (Figure 4a).

### Genes evolving rapidly in *T. dactyloides* also experienced selection in temperate adapted maize

Previous studies have identified a large set of maize genes which were targets of artificial selection during the process of adaptation to temperate climates [20]. We hypothesized that the more ancient process of the expansion of *T. dactyloides* into temperate climates may have targeted some of the same genes targeted by artificial selection during the introduction of maize into temperate climates. Tripsacum genes were divided into those where the maize ortholog was identified as likely under selection during the adaption of maize to temperate climates and those where the maize ortholog did not show evidence of being under selection during this process. As Ka/Ks ratios can vary widely across different genes as a result of factors including expression level, gene functional category, and location relative to centromeres [21] all *T. dactyloides* Ka/Ks values were normalized relative to sorghum, a closely related species that is still primarily adapted to tropical latitudes. Genes under selection during the development of temperate maize lines showed significant increases in Ka/Ks values in temperate adapted *T. dactyloides* relative to tropical adapted sorghum (log transformed t-test *p*-value = 0.027, Wilcoxon signed-rank test (WST) *p*-value = 0.018) (Figure 3B) [22]. This observation remained significant when using the median Ka/Ks value from orthologs in three different tropically adapted grass species (rice, oropetium and sorghum) (log transformed t-test *p*-value = 0.038, WST *p*-value = 0.029).

While the Hufford dataset consisted primarily of temperate elite lines and tropical landraces, it did also include a number of elite tropical lines and temperate landraces. A second dataset consisting of 47 high confidence tropical and subtropical maize lines and 46 high confidence temperate lines from maize HapMap3 [23], provided approximately equivalent results (Table S8). Normalized Ka/Ks values of *T. dactyloides* genes orthologous to the 10% of maize genes with the highest XP-CLR scores were not significant higher than the background (p-value = 0.24 in log transformed t-test, 0.19 in WST). However the normalized Ka/Ks value of top 5% and top 1% genes were significantly higher than background genes (top 5%: *p*-value = 0.032 in log transformed t-test, 0.018 in WST; top 1%: *p*-value = 0.028 in log transformed t-test, 0.008 in WST). The increase in significance increased at more stringent cut offs – even as the total number of data points decreases – may indicate that the overlap comes from only a subset of the genes under the strongest selection between tropical and temperate lines in maize and in the temperate adapted *T. dactyloides* relative to tropical-latitude-adapted related species.

## Discussion

The potential for data from *Tripsacum* to aid in both basic biological research and applied plant breeding in maize has long been discussed [9, 10]. However, as of December 2017, a total of only 611 published nucleotide sequences existed for the entire *Tripsacum* genus, including 565 for *T. dactyloides*, 12 for *T. andersonii*, and 34 for all other named taxa within the genus. Here we have generated a set of 24,616 full length *T. dactyloides* cDNA sequences covering 22.4%, 31.5% and 60.2% of the annotated, expressed, and syntenically conserved gene space of maize respectively. This larger scale transcriptome resource enables a number of comparative analyses of *Zea* and *Tripsacum* not previously feasible.

Significant evolutionary rate heterogeneity exists among extant grass species. Previously, variation in the rate of divergence between homeologous gene pairs generated during the rho polyploidy [24] in different grass species was employed to detect variation in synonymous substitution rates [25]. However, this approach, which relies on pairwise comparisons between species, provides aggregate estimates for each lineage across the 70–96 million years since the rho WGD [24, 25]. Utilizing known phylogenetic relationships across relatively large numbers of grass species with sequenced genomes or significant genome resources and fitting rates of synonymous and nonsynonymous substitutions for each branch separately [26] demonstrated that even comparing sister genera *(Zea* and *Tripsacum)*, maize exhibits significantly more rapid accumulation of synonymous substitutions. One potential explanation is differences in life cycle. Many *Zea* species are annuals [4] while more than 13 *Tripsacum* species are perennials [5] (Table 1) and several analyses have suggested that synonymous substitutions accumulate more rapidly in annual species [27, 28]. Another potential explanation is that the difference in the rate of molecular evolution between maize and *T. dactyloides* may reflect the difference in native range between these genera, as species native to tropical regions have been shown to accumulate nucleotide substitutions up to twice as rapidly as temperate species [29].

Genes are generally considered to show evidence of positive selection if the frequency of nonsynonymous substitutions is significantly higher than that of synonymous substitutions. However, if positive selection is assumed to be episodic rather than constant, elevated Ka/Ks ratios which are less than one can reflect a mixture of positive selection and purifying selection, relaxation of purifying selection, or statistical noise. Episodic positive selection is harder to detect on longer branches where the proportion of evolutionary time a gene spends under positive selection decreases relative to the time spent under purifying selection. Background ratios of Ka/Ks can also vary significantly between different genes reflecting differences in chromosome environment, expression level, function, and the presence or absence of different types of duplicate gene copies [21]. As shown in Figure 2C, the frequency of extreme Ka/Ks ratios decreases as the overall branch length increases. The inclusion of *T. dactyloides* breaks up the long branch between maize and sorghum, permitting the identification of genes experiencing either an interval of positive selection alongside ongoing purifying selection or a relaxation of purifying selection. It must be emphasized that the analysis presented here cannot distinguish between true positives – genes showing elevated rates of protein sequence evolution as a result of positive selection or relaxed selection – and false positives – statistical noise – on a single gene level. Instead the focus must be on the differences observed between the functional classes or pathways of genes which exhibited higher rates of protein sequence evolution in maize and *T. dactyloides*. Here we found that genes involved in phospholipid metabolism and stress response both tended to be experiencing higher rates of protein sequence evolution in *T. dactyloides* than in maize or in the other grass species tested.

Recent studies of several instances of parallel selection have reported that it often acts on largely unlinked sets of genes at a molecular level [30, 31, 32]. This suggests that there are many different molecular mechanisms which can be employed to achieve the same phenotypic changes, and as a result the same genes will rarely be targeted in independent instances of selection for the same traits. However, the evidence presented above suggests that as the lineage leading to *T. dactyloides* expanded into temperate environments millions of years ago natural selection targeted some of the same genetic loci which would later be targets of artificial selection as the cultivation of maize spread from the center of domestication in Mexico into more temperate regions of North America. This overlap between targets of natural and artificial selection for adaptation to the same environment in sister genera also indicates that genetic changes in crop-wild relatives associated with adaptation to new environments may be useful guides for identifying genetic targets for breeding efforts aimed at adapting crops to a changing climate.

One specific difference between the native environment of wild *Zea* and *Tripsacum* species is that many *Tripsacum* species grow in areas where they are exposed to cold and freezing temperatures. Unlike maize, *T. dactyloides* can survive prolonged cold and freezing temperatures and successfully overwinter. The identification of accelerated protein sequence evolution among genes involved in phospholipid metabolism provides a plausible candidate mechanism for the increased cold and freezing tolerance of *T. dactyloides* relative to maize. While widely grown in temperate regions over the summer, maize remains sensitive to cold. Maize varieties with the ability to be planted significantly earlier, or in the extreme case to overwinter, have the potential to intercept a greater proportion of total annual solar radiation increasing maximum potential yields [33, 11]. This study illustrates how studying the genetic mechanisms responsible for crop-wild relative adaptation to particular climates may guide breeding and genome engineering efforts to adapt crops to a changing climate.

## Methods

### Plant materials and RNA preparation

*T. dactyloides* seeds were collected from wild growing plants located in Eastern Nebraska (USA, GPS coordinates: 41.057836, −96.639844). Seeds with brown cupules were selected. For each seed the cupule was removed, followed by a cold treatment at 4°Cfor at least two days, resulting in germination rates between 30% and 50%. A single plant (ID #-1) was selected for transcriptome sequencing. Young leaves were sampled one month after germination. Stem and root were sampled three months after germination. Harvested tissue samples were rinsed with cold distilled water and then immediately frozen in liquid N_2_. Total RNA was extracted from each tissue separately by manually grinding each sample in liquid N_2_, adding TriPure isolation reagent (Roche Life Science, catalog number #11667157001), followed by separating phase using chloroform, precipitating RNA using isopropanol and washing the RNA pellet using 75% ethanol. The air-dried RNA samples were dissolved in DEPC-treated water. RNA quantity and quality were assessed uisng a NanoDrop 1000 spectrophotometer and electrophoresis on a 1% agarose gel respectively.

### Single-molecule sequencing and isoform detection

Equal quantities of total RNA from each sample were pooled prior to library construction and the combined sample was shipped to the Duke Center for Genomic and Computational Biology (GCB), Duke University, USA for sequencing. Three size-fractionated libraries (1–2 kb, 2–3 kb, and 3–6 kb) were constructed and sequenced separately on the PacBio RS II. Each libraries was sequenced using 2 SMRT cells. Raw reads data was analyzed through running the Iso-Seq pipeline included in the SMRT-Analysis software package (SMRT Pipe v2.3.0, https://github.com/PacificBiosciences/cDNA_primer) (for details see Supplemental Methods).

### Substitution rate estimation and selection analyses

Codon based alignments were generated using ParaAT2.0 [34] for sets of orthologous genes identified using a dataset of syntenic orthologous genes identified across six grass species with sequenced genomes (maize V3 [35], sorghum v3.1 [36], setaria v2.2 [37], oropetium v1.0 [38], rice v7 [39] and brachypodium v3.1 [40]) [17], plus the maize/tripsacum orthologous relationships defined above. Synonymous nucleotide substitution rates (Ks) were calculated by using the codeml maximum-likelihood method (runmode = −2, CodonFreq = 2) implemented within PAML [26] and the known phylogenetic relationships of the seven species. The divergence time (T) between maize and *T. dactyloides* was estimated following the formula T = Ks/2*μ*[41] (for more details see Supplemental Methods).

### Genome-wide scan for selection between tropical/temperate maize subpopulations

A cross-population composite likelihood approach XP-CLR [42] (updated by Hufford et al.[20] to incorporate missing data), based on the allele frequency differentiation between purely tropical/temperate maize subpopulations was used to identify genes likely to have been targets of selection during the process of adaption from tropical to temperate climates. 47 tropical and subtropical lines (TSS) and 46 temperate lines (NSS and SS) were selected from the Hapmap3 dataset [23], based on having a the probability value of membership into either of these groups greater than than 0.8 (Table S8) [43]. Based on the recombination rates measured using high density genetic maps in maize [44], XP-CLR was run with the following parameters: sliding window size of 0.05 cM; a fixed number of SNPs per window of 100; downweighting of SNPs in high LD (r2 > 0.75); and a set of grid points as the putative selected allele positions were placed with a spacing of 1 kb across the whole genome. Each gene was assigned the maximum XP-CLR values found within the region 5 kb up- and downstream of the gene’s annotation transcription start and transcription stop site.

### Profiling of lipid and transcriptional responses to cold

Maize, sorghum, and *T. dactyloides* seedlings were grown under 13 hours/11 hours 29 °C /23 °C day/night and 60% relative humidity in a growth chamber at University of Nebraska-Lincoln’s Beadle Center facility. At the three leaf stage, one half of the seedlings were moved to a second growth chamber maintained at a constant 6 °C temperature. The initiation of cold stress was timed to coincide with the end of daylight illumination. For RNA, *T. dactyloides* seedlings were harvested from both control and cold stressed conditions at 1, 3, 6, and 24 hours after the initiation of cold stress treatment. For lipid characterization, seedlings were harvested from control and cold stressed plants 24 hours after the initiation of cold stress. Raw fastq sequences for maize and sorghum cold stress treatment are those published [45]. All data was realigned and reanalyzed using Trimmomatic [46], GSNAP [47] and HTSeq [48]. Differentially expressed genes (DEGs) and differentially regulated orthologs (DROs) were identified from read count data using DESeq2 [49], as described in [45].

Lipid composition determined essentially as described [50] (see Supplemental Methods for details).

### Data availability

Raw PacBio sequence data has been deposited in the NCBI SRA under project PRJNA471728. Raw Illumina RNA-seq data has been deposited in the NCBI SRA under project PRJNA471735. A fasta file with the set of non-redundant *T. dactyloides* isoforms employed in this study is available at Zenodo with the identifier http://dx.doi.org/10.5281/zenodo.841005.

## Acknowledgements

We thank Dr. Christy Gault (Cornell University) for sharing her protocol for germinating *T. dactyloides* seeds. This work was supported by the National Science Foundation under Grant No.OIA-1557417 to JCS, USDA NIFA award 2016-67013-24613 to RLR and JCS, a Science Foundation of Xichang University awarded to LY and a China Scholarship Council fellowship awarded to XL.

## Author contributions

JCS, RLR, LY and XL conceived the project and designed the studies; OR identified and collected the plant material used in this study; LY, YZ, and SM performed the experiments; LY, XL, SKKR, and XD analyzed the data; LY, SKKR, and JCS wrote the paper. All authors reviewed and approved the final manuscript.

## Supplemental Information (SI)

### Supplemental Results

A single *Tripsacum dactyloides* plant grown from seed collected from the wild in eastern Nebraska (USA) was used as the donor for all RNA samples. RNA extracted from three tissues (root, leaf, and stem) was used to construct three size fractionated libraries (1–2, 2–3, and 3–6 kb) which were sequenced using a PacBio RS II yielding a total of 532,071 Read Of Inserts (ROIs). The SMRT Pipe v2.3.0 classified more than half (267,186, 50.2%) of the ROIs as full-length and non-chimeric (FLNC) transcripts based on the presence of 5’-, 3’-cDNA primers and polyA tails. Each size-fractionated library had expected average length of FLNC transcripts of 1,364 bp, 2,272 bp, and 3,323 bp, with the average length of total FLNC transcripts of 2,354 bp, ranging from 300 to 29,209 bp Table S1). ICE and Quiver processing of FLNC transcripts produced a total of 64,362 high quality (HQ) consensus transcript sequences with an estimated consensus base call accuracy ≥ 99% (Figure S4).

Final consensus tripsacum sequences were mapped to the maize reference genome (RefGen_v3) using GMAP. Consistent with previously reported low overall rate of divergence in gene content and gene sequence between maize and tripsacum [9], 98.04% (63,103 out of 64,362 HQ consensus sequences) could be confidently mapped to the maize reference genome. Pbtranscript-TOFU was used to collapse the consensus sequences into 24,616 unique isoforms, in which differences in the 5’ end of the first exon were considered redundant, and otherwise identical isoforms were merged (see Methods). This final set of unique tripsacum isoforms mapped to a total of 13,089 maize genes, including 7,633 maize genes represented by a single tripsacum transcript and 5,456 maize genes represented by two or more transcripts. Among maize genes to which two or more tripsacum isoforms were mapped the average was 3.1 isoforms per gene. Eighty-four maize genes were represented by more than 10 or more tripsacum isoforms, and the single maize gene represented by the most isoforms was GRMZM2G306345, which encodes a pyruvate, phosphate dikinase (PPDK) protein involved in the fixation of carbon dioxide as part of the C4 photosynthetic pathway, with 83 identified tripsacum isoforms (Table S2). A set of 249 confident lncRNAs were identified among the final tripsacum consensus sequences (See Methods) (Figure S6G). With an average length of 1.45 kb (ranging from 0.51 to 3.5 kb), the distribution of lncRNA sequences is notably larger than the average length of the maize lncRNAs identified using pacbio isoseq 0.67 kb (ranging from 0.2–6.6 kb) (Figure S5) [56]. Only 17 of these 249 lncRNAs exhibited high sequence similarity (identity > 80%) with lncRNA sequences identified in maize.

In total, 12,826 out of 13,089 tripsacum transcripts mapped to 14,401 annotated maize gene models (Figure S6A,B). In some cases a single consensus tripsacum sequence spanned two or more maize gene models (Figure S6). Maize genes were sorted based on their expression levels using an existing short read RNA-seq dataset from maize seedling tissue [45]. Of the 13,089 most highly expressed maize genes, 8,191 (62.6%) were aligned with at least one tripsacum transcript indicating that the expression level of a gene in maize is a relatively good predictor of how likely that gene was to be captured in the tripsacum isoseq data. Because genes located in the chromosome arms of maize tend to exhibit higher levels of expression than genes in pericentromeric regions, this sample of tripsacum genes is likely depleted in the types of genes which are over represented in pericentromic regions. Figure S6C and D illustrate the comparative densities of highly expressed maize genes aligned to tripsacum isoseq transcripts across the 10 chromosomes of maize. Tripsacum transcript density was correlated with the density of maize highly expressed genes (Spearman correlation coefficient r = 0.855, *p <* 2.2e-16). Among the 12,826 tripsacum transcripts mapped to maize gene models, 11,910 were unique one-to-one mappings. Data on conserved maize-sorghum orthologous gene pairs was used to increase the confidence of these mappings (Figure S6E), resulting in a final set of 9,112 putative sorghum-maize-tripsacum orthologous gene groups (available on https://figshare.com/s/6d55867b09e014eb7aed, Figure S6F). These gene pairs were grouped into three categories “one-to-one”, single tripsacum/maize orthologs without duplication (5,641 gene pairs); “one-to-two”, a single tripsacum gene mapped to one copy of a homeologous maize gene pair (1,964 gene triplet); “two-to-two”, unique tripsacum sequences mapped to each copy of a homeologous maize gene pair (1,507 gene quartets, Figure S6H, Figure S7).

A total of 223 transcripts aligned to the maize genome in locations where no maize gene model was present. After additional validation and QC (see Methods), 94 of these cases appeared to be unannotated maize genes which were supported by both an aligned tripsacum transcript and short read RNA-seq data in maize, and another 102 cases appeared to be genomic sequences conserved between maize and tripsacum which were transcribed in tripsacum but lacked expression evidence in maize. More than two thirds of the 94 potentially unannotated maize genes could be confidently aligned to annotated coding sequences in the reference genomes of sorghum (63%) or setaria (66%). In contrast, the sequences present in both the tripsacum and maize genomes but expressed only in tripsacum were much less likely to be present in outgroup species (sorghum (25%) and setaria (30%)), suggesting the transcription of these genomic sequences may be a comparatively recent feature, potentially unique to the tripsacum lineage.

Consistent with observations from other plant systems [57, 58, 59], the most common single AS variants in tripsacum were – in descending order of frequency – intron retention (IntronR), exon skipping (ExonS), alternative donor (AltD), alternative acceptor (AltA), and mutually exclusive exons (MXEs). The remaining isoforms incorporated two or more types of changes and were classified as Complex AS (CompAS) (Figure S10). While a comparable maize dataset generated using the same long read technology contained more total splicing events, likely as a result of approx 4.5x greater sequencing depth [56], the overall proportions of AS events belonging to different categories in maize and tripsacum were more similar to each other than either was to sorghum (Figure S10). Shifting from overall frequency to the conservation of specific splicing events – defined as identical AS codes at the same physical positions when tripsacum and maize full length transcripts are aligned to the maize reference genome – a total of 4,324 (35.3%) tripsacum AS events associated with 1,065 genes were also identified in maize (Figure S10) and more than two third (656, 61.6%) of the conserved AS genes were observed in orthologous gene groups while 409 genes were *Zea-Tripsacum* linage-specific in *Tripsacinae*.

Of the 1,542 gene quartets where a single gene in sorghum is co-orthologous to two maize genes and each maize gene is orthologous to a single tripsacum gene, 212 genes exhibited alternative splicing for both tripsacum genes and 409 genes exhibited alternative splicing for both maize genes. In 52.06% of of gene pairs in tripsacum the pattern of splicing was conserved between homeologous gene pairs as well as in 77.8% of maize homeologous gene pairs. Changes in patterns of splicing which could be more parsimoniously explained by differences in *trans-* regulation of AS were observed substantially substantially for frequently than were changes in changes in patterns of splicing which could be more parsimoniously explained by differences in *cis-* regulation (Figure S11).

### Supplemental Methods

#### Creation of high quality consensus transcript sequences

First, reads of insert consensus sequence (ROIs, previously known as circular consensus sequences, CSCs) were identified using the Reads Of Insert protocol, one of the submodules of the Iso-Seq pipeline. The minimum required number of full passes was set to 0 and predicted consensus accuracy was set to 75%. Raw sequences that passed this step were considered to be the ROIs.

Second, the classify submodule was used to automatically determine which ROIs were full-length. ROIs were considered to be full length if both the 5’- and 3’-cDNA primers as well as a poly(A) tail signal preceding the 3’-primer were detected. The classify submodule also separated chimeric reads from non-chimeric reads through detecting the presence of SMRTbell adapters in the middle of sequences, producing a set of full-length non-chimeric (FLNC) reads.

Third, the isoform-level clustering algorithm ICE (Iterative Clustering for Error Correction), which uses only full length reads, followed by Quiver which polished FLNC consensus sequences from ICE using non-FL reads were used to improve the accuracy of FLNC consensus sequences. These steps resulted in a set of high quality polished consensus with > 99% post-correction accuracy.

Fourth, HQ polished consensus reads were mapped to the maize reference genome (B73_RefGen_v3) using GMAP and redundant mapped transcripts were merged using using the collapse_isoforms_by_sam.py script from the pbtranscript-ToFU package (https://github.com/PacificBiosciences/cDNA_primer/wiki/tofu-Tutorial-(optional).-Removing-redundant-transcripts/) with the parameter settings of min-coverage = 85% and min-identity =82%. Isoforms differing only at the 5’-sites within the first exon were considered to be redundant and collapsed.

#### Ka/Ks Analysis Details

Branch-specific Ka/Ks ratios with self-built phylogenetic trees were calculated using codeml program (runmode = 0). The resulting codeml data including Ka, Ks and Ka/Ks for the genes of each branch were obtained and genes with Ka > 0.5, Ks > 2 and Ka/Ks > 2 were discarded. Two models in codeml were used, one required a constant value for ω across all branches (model 0), and the other allowed heterogenous rates on each branch of the phylogeny (model 1). A likelihood ratio test was used to compare likelihood values under model 0 and model 1 to test whether significant variation in Ka/Ks ratios between different branches was present [60]. For each set of orthologous genes the log likelihood values under two models (lnL1 for the alternative and lnL0 for the null model) were obtained. From these values, the LRT was computed using 2 × (lnL1-lnL0). χ^2^ curve with degree of freedom = 1 was used to calculate a *p*-value for this LRT. Multiple testing was performed to correct these *p*-values by applying the false discovery rate method (FDR) adjusted in R [61]. A gene was considered to be experiencing accelerated rates of protein evolution in a given lineage if the highest Ka/Ks ratio was detected in the branch leading to that species and the FDR-adjusted *p*-value for the comparison of the constant ω and heterogeneous models was < 0.05.

#### Use of statistical tests in comparisons of Ka/Ks ratios between different populations of genes

Like gene expression fold-change data, Ka/Ks ratios and ratios of Ka/Ks ratios exhibit non-normal distributions. Applying a log transformation to either Ka/Ks ratios or ratios of ratios produces roughly normal distributions, as observed for gene expression fold change data [22]. This method was employed to conduct two comparisons using the independent-samples t-test package within R. In addition, a non-parametric test in R, Wilcoxon signed-ranked tests, were also used test whether Ka/Ks ratios differ significantly between gene groups. In the first test, the distribution of ratio of ratio values between *T. dactyloides* and sorghum Ka/Ks values was compared between genes identified as likely to be under selection in a comparison of tropical and temperate maize lines [20, 32] and genes not identified as likely to be under selection in the same comparison. The second test compared Ka/Ks ratios for genes annotated as involved in lipid metabolism to the population of genes not involves in phospholipid metabolism in *T. dactyloides* and maize, respectively, as well as Ka/Ks ratios between *T. dactyloides* and maize.

#### Lipid Profiling Details

Briefly, lipid extraction was performed by immediately immersing fresh tissue samples in ice-cold 2:1:0.1 (v/v/v) methanol: chloroform: formic acid and bead beating them for 2 minutes until the tissue was completely disrupted, then phase-partitioning by addition of 0.2M phosphoric acid, 1 M potassium chloride. The organic phase was removed (and stored at −20 degrees Celsius until) derivatization. 2D-thin Layer Chromatography (TLC) was used to separate lipids by their headgroup properties using the following solvent systems, 130:50:10 (v/v/v) Chloroform: Methanol: Ammonia hydroxide in the first dimension and 85:12.5:12.5:4 (v/v/v/v) Chloroform: Methanol: acetic acid: water in the second dimension. Lipids were identified by with respect to standards purchased from Avanti Polar Lipids. Quantitative data was obtained by exposure to iodine vapor and compared with a standard mixture. Each target lipid was scraped from the TLC plate and derivatized to fatty acyl methylesters (FAME Reaction) and the resulting profile of FAME was determined by gas chromatography (GC).

#### Identification of LncRNAs

The protein coding-potential of individual *T. dactyloides* transcripts was assessed using CPAT (Coding Potential Assessment Tool) [62], which employs a logistic regression model built with four sequence features as predictor variables: open reading frame (ORF) size, ORF coverage, Fickett testcode statistic, and hexamer usage bias. CPAT was trained using 4,900 high-confidence lncRNAs transcripts identified as part of Gramene 52 release (http://www.gramene.org/release-notes-52/) [63] and an equal number of known protein-coding transcripts randomly subsampled from the RefGen_v3 annotation 6a to measure the prediction performance of this logic model in maize. The accuracy of the trained CPAT model was assessed by quantifying six parameters using 10-fold cross validation with the maize training dataset: sensitivity (TPR), specificity (1-FPR), accuracy (ACC) and precision (PPV) under the receiver operating characteristic (ROC) curve and precision-recall (PR) curve. Based on there parameters, a probability threshold of 0.425 was identified as providing the best trade off between specificity and sensitivity for identifying lncRNA sequences.

4,095 candidate non-coding RNAs with a length greater than > 200 bp were predicted in CPAT. ORF prediction for non-coding candidate of *T. dactyloides* transcripts was performed using TransDecoder [64] and 2,509 transcripts encoding ORFs longer than 100 amino acids were removed from the set of putative lncRNAs. The remaining lncRNAs were aligned to the NCBI-nr database using BLASTX and sequences showing similarity to existing protein sequencing in the database (e-value 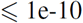) were removed from the set of putative lncRNAs.

#### Identification of orthologs

Maize-tripsacum orthologs were defined based on the following criteria. Firstly, the tripsacum sequence must have been uniquely mapped to the target maize gene using GMAP. Secondly, the tripsacum gene must have been uniquely mapped to a single sorghum gene using BLASTN. Thirdly, the maize and sorghum genes must be syntenic orthologs of each other using a previously published dataset [17, 45]. As a result of the maize WGD shared by maize and *T. dactyloides*, in some cases multiple *T. dactyloides* sequences mapped to unique maize genes on different maize subgenomes, but to the single shared co-ortholog of these two maize genes in sorghum.

#### BLAST analyses for Isoseq data

Isoforms which failed to map to the maize genome aligned to different NCBI refSeq databases using BLASTN with the following parameters: min-coverage = 50%, min-identity = 70%, max_target_seqs = 5 and e-value 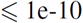. Transcripts were considered to originate from ancestral grass genes not present in the B73 reference genome if none of the top 5 blast hits were to maize sequences, but did include sequences isolated from other grass species.

*T. dactyloides* sequences which aligned to regions of the maize genome not annotated as genes were also aligned to the full set of annotated maize gene CDS sequences using BLASTN with parameters: min-coverage=80%, min-identity=85%, max_target_seqs=1 and e-value 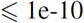. Transcripts which did not align to any maize gene models even with these relaxed criteria were then manually proofed for expression evidence in maize using IGV and a set of Illumina short read RNA-seq data from a wide range of maize tissues [65].

#### Detection of alternative splicing (AS) events

AS analysis was conducted using Astalavista-4.0 (Alternative Splicing Trascriptional Landscape Visualization Too) [66], transforming identified place isoforms into a set of defined AS codes based on their location within the gene and the type of splicing observed. Among the multiple maize genes with multiple aligned *T. dactyloides* isoforms, in 3,566 cases one or more AS events (12,260 in total) was identified between the multiple aligned isoforms, while in the 1,890 remaining cases differences between isoforms were the result of either variation in transcription start sites (VSTs), or additional 3’ exons (DSPs). Each AS site is assigned a number according to its relative position in the event and a symbol depending on its type. In addition to the main five single AS types such as Intron retention (IR), Exon skipping (ES), Alternative donor (AD), alternative acceptor (AA) and Mutually exclusive exons (MXEs), other AS events in which multiple types of AS are present between two isoforms were counted as complicated AS. The maize data used to compare AS events between maize and *T. dactyloides* was extracted from the published AS dataset generated by Wang et al [56].

**Table S1.** Summary of sequence data produced.

**Table S2.** Number of maize genes with one or more identified tripsacum isoforms.

**Table S3.** Patial list of conserved grass genes present in tripsacum but not in maize reference genome.

**Table S4.** Genes experiencing accelerated selections in *T. dactyloides* related to stress response.

**Table S5.** GO terms associated with Maize-Tripsacum and Sorghum Tripsacum DROs after 3, 6, 12 and 24 hours post cold stress

**Table S6.** GO terms associated with up-regulated and down-regulated Maize-Tripsacum and Sorghum Tripsacum DROs after 6 hours post cold stress

**Table S7.** ANOVA table for lipid desaturation levels in maize, sorghum and *T. dactyloides* at room temperature and 24 hours after cold stress.

**Table S8.** List of purely temperate and tropical maize lines from HapMap3.

## Supplemental Figures

**Figure S1.**
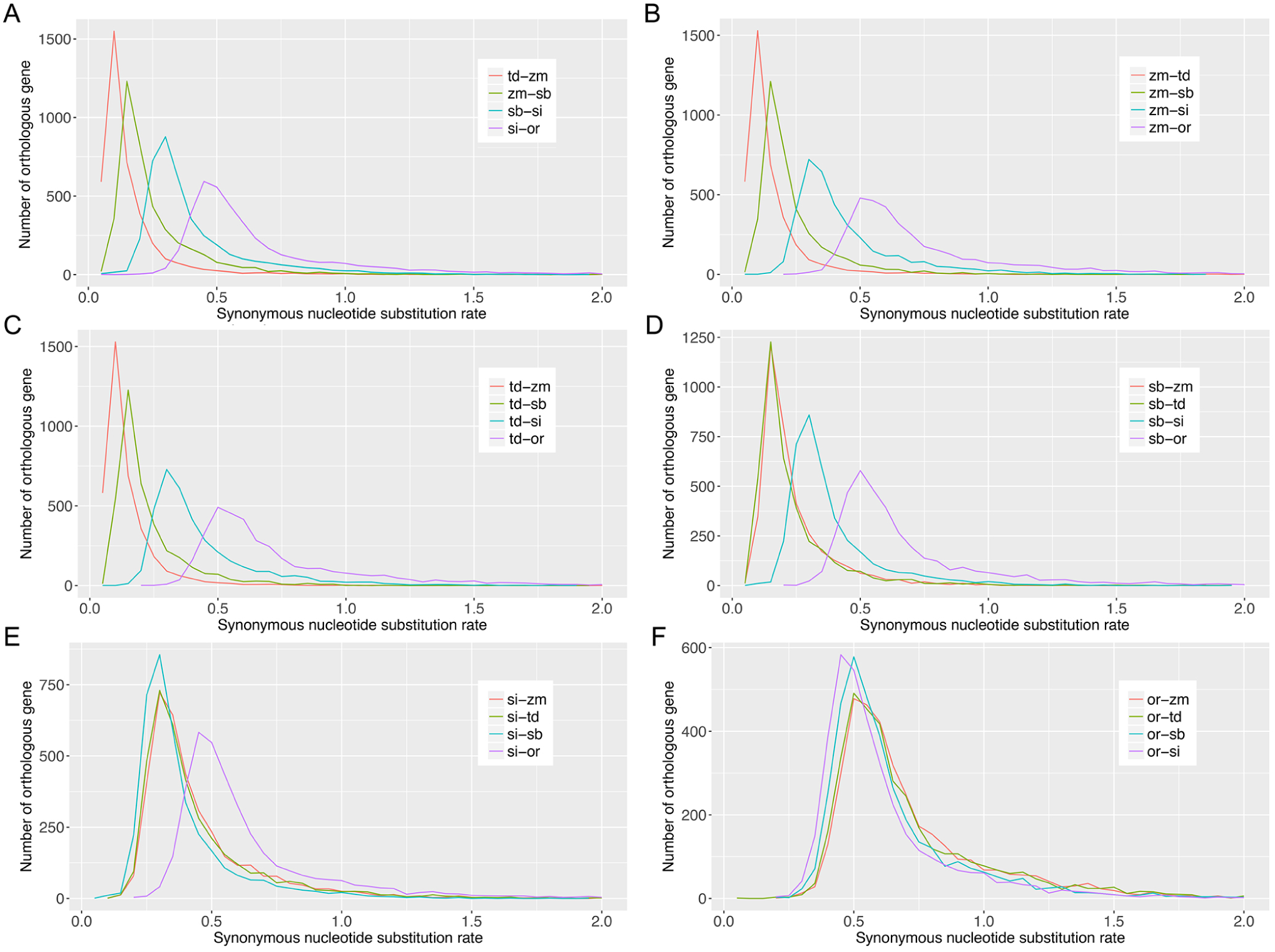
Ks distribution of orthologous gene pairs between each two of species pair-wisely (bin size = 0.05). (A) divergence between tripsacum and maize (td-zm), maize and sorghum (zm-sb), sorghum and setaria (sb-si), setaria and oropetium (si-or), their divergence were shown by the peak of each pair. (B) divergence of maize with other species. (C) divergence of tripsacum with other species. (D) divergence of sorghum with other species. (E) divergence of setaria with other species. (F) divergence of oropetium with other species.

**Figure S2.**
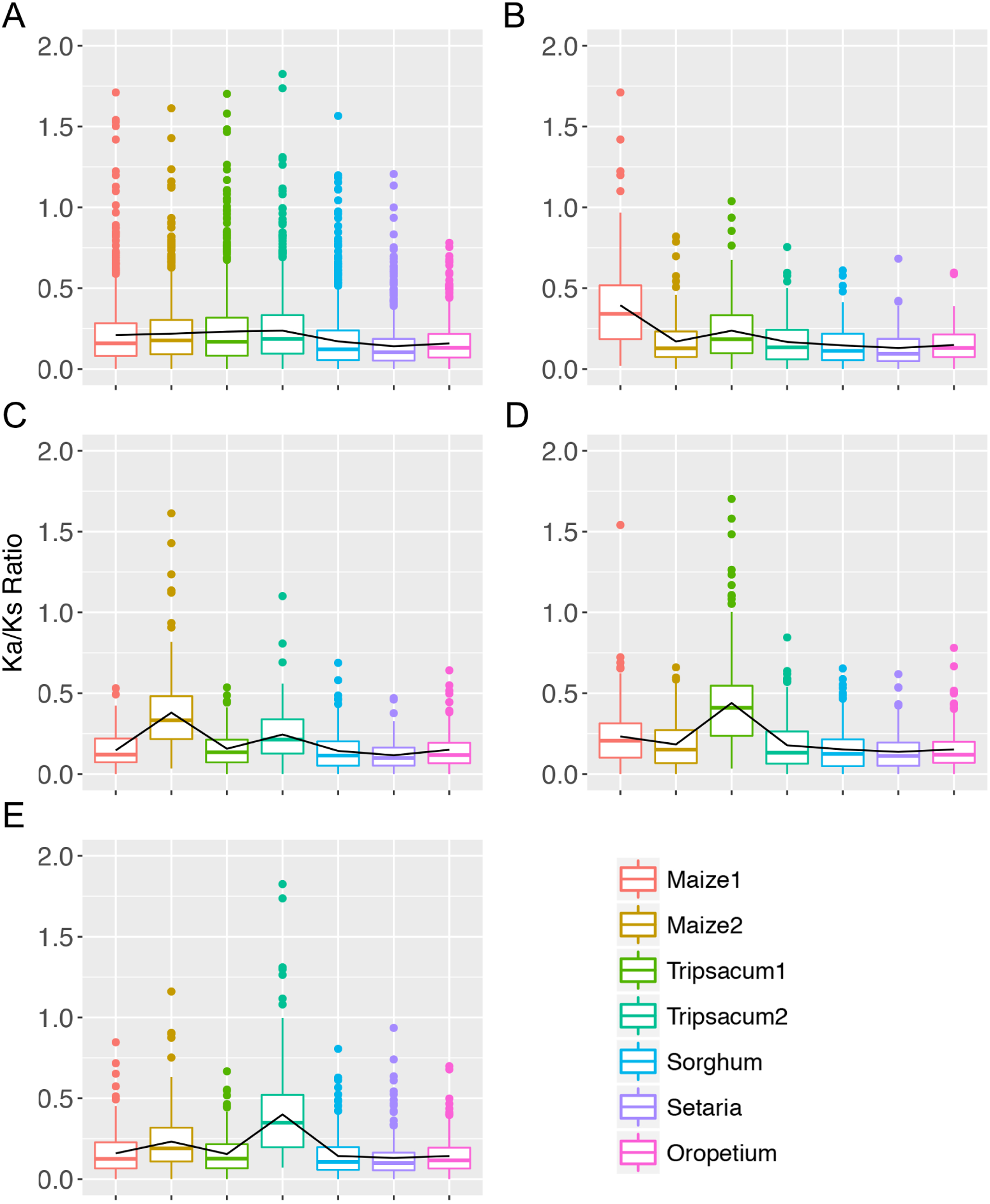
Distribution of species-specific Ka/Ks ratios including the homeologous gene quartets in both tripsacum and maize shared the WGD. (A) distribution of Ka/Ks ratios in orthologous genes sets. (B) increased Ka/Ks ratios in maize1. (C) increased Ka/Ks ratios in maize2. (D) increased Ka/Ks ratios in tripsacum1. (E) increased Ka/Ks ratios in tripsacum2.

**Figure S3.**
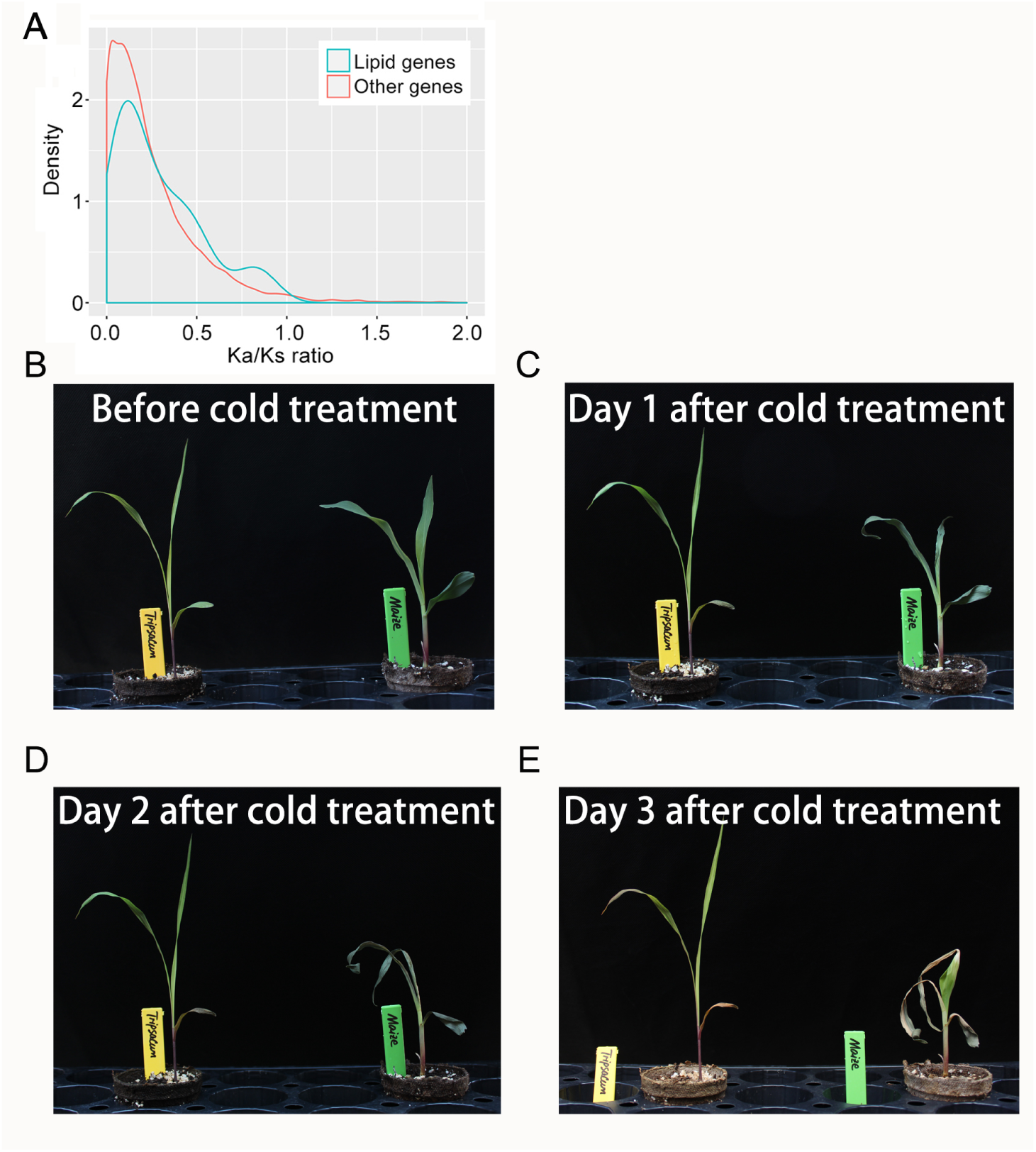
(A) Ka/Ks distributions in tripsacum between lipid genes and other functional genes. (B-E) Phenotype changes between tripsacum (left) and maize (right) in continuous three days after cold treatment at 4 °C for five days.

**Figure S4.**
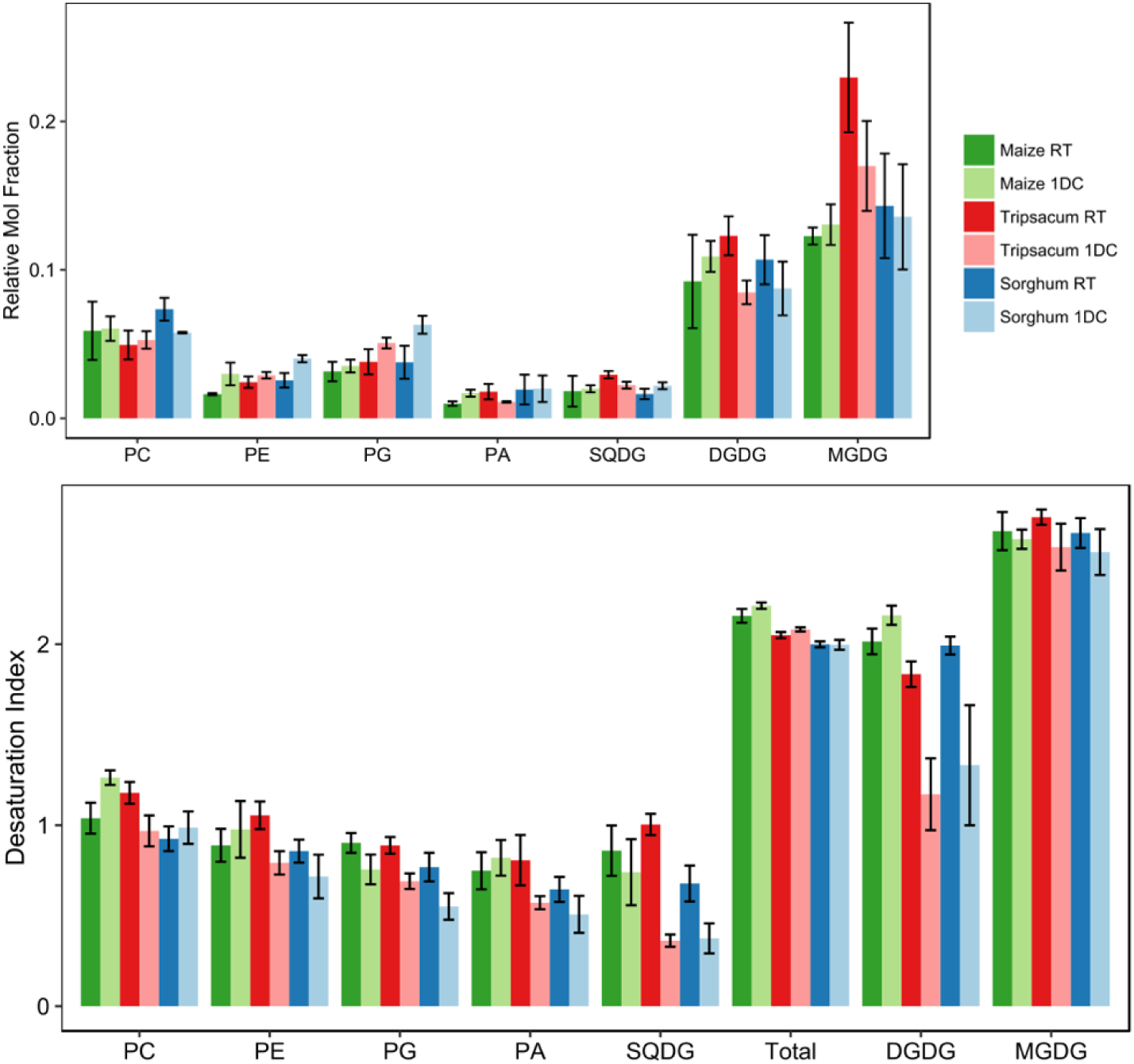
Changes in abundance (top) and desaturation (bottom) of specific lipid types in seedlings of maize, *T. dactyloides*, and sorghum under control conditions (RT) and after exposure to 24 hours of cold stress (1DC).

**Figure S5.**
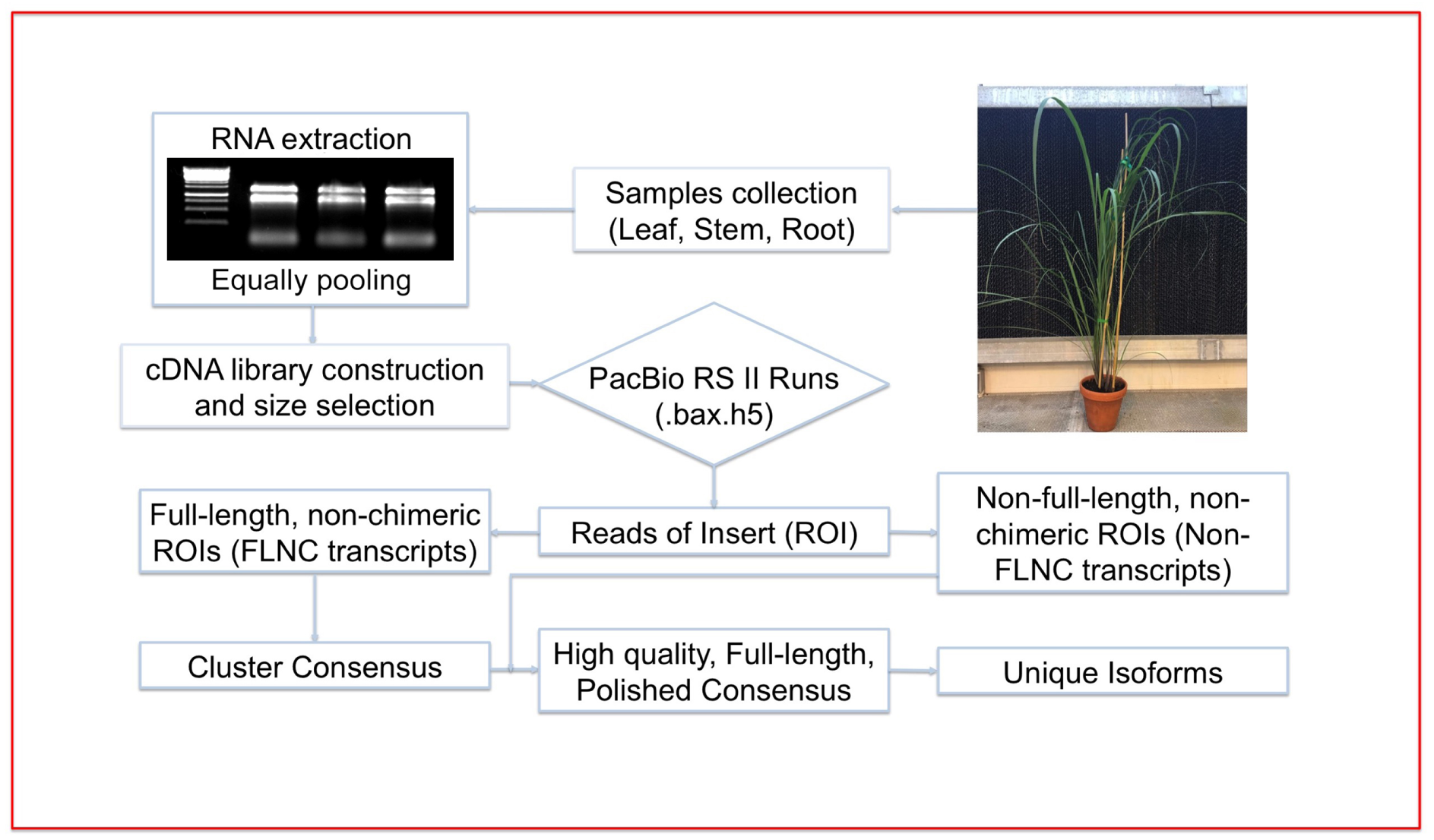
Workflow of Iso-Seq bioinformaics analysis for tripsacum using the SMRT-Analysis software package (SMRT Pipe v2.3.0).

**Figure S6.**
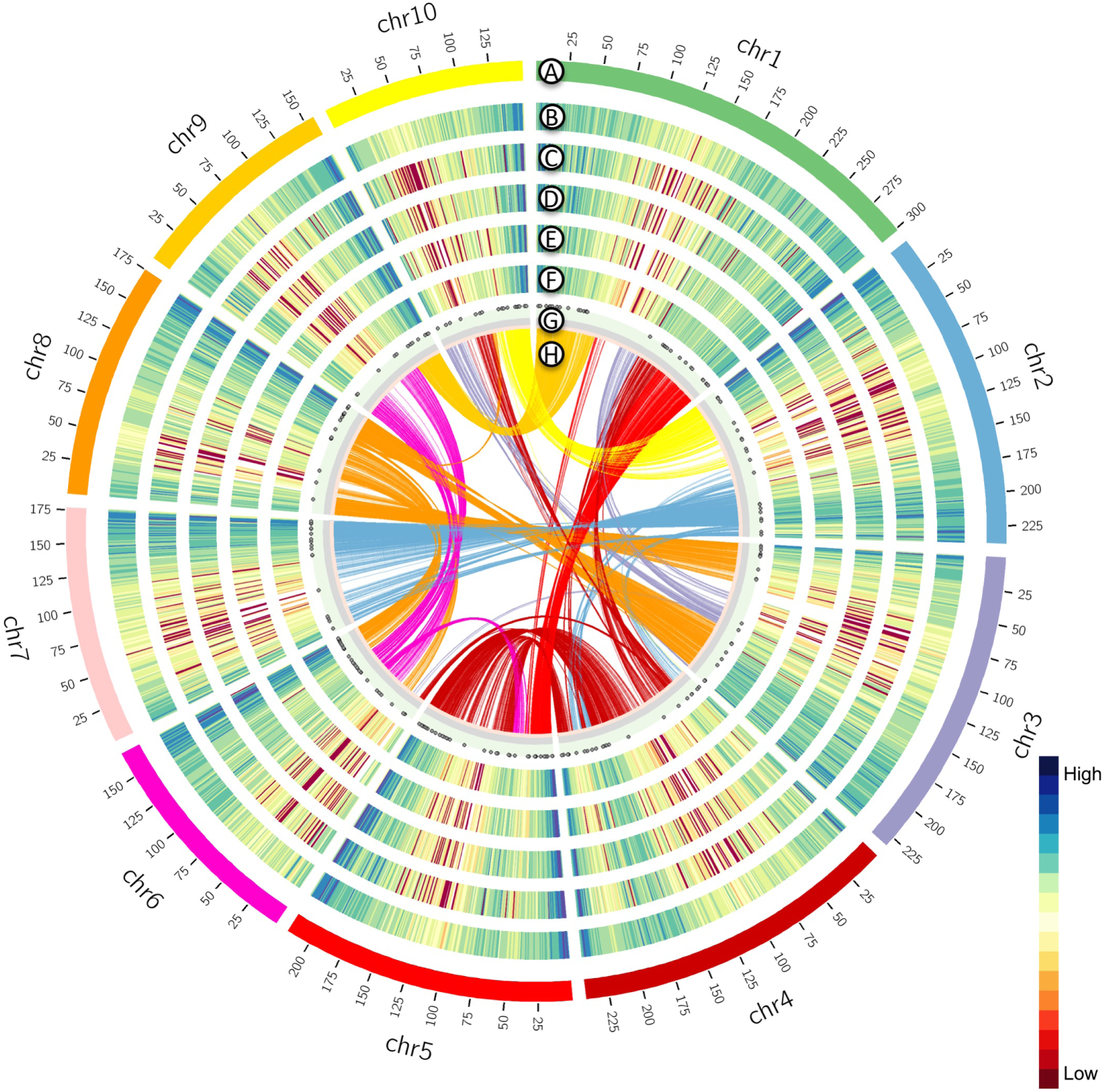
Tripsacum Iso-seq data mapped onto the maize reference genome (RefGen_v3). (A) 10 chromosomes of maize. (B) Maize gene density in each chromosome. (C) Density of highly expressed maize genes (FPKM > 4.15). (D) Tripsacum transcript density in each chromosome. (E) Density of syntenic gene pairs between maize and sorghum. (F) Density of sorghum-maize-tripsacum gene pairs. (G) Distribution of Long non-coding RNA (IncRNA) in tripsacum. (H) Distributions of homeologous genes pairs following the whole genome duplication which is shared by both tripsacum and maize.

**Figure S7.**
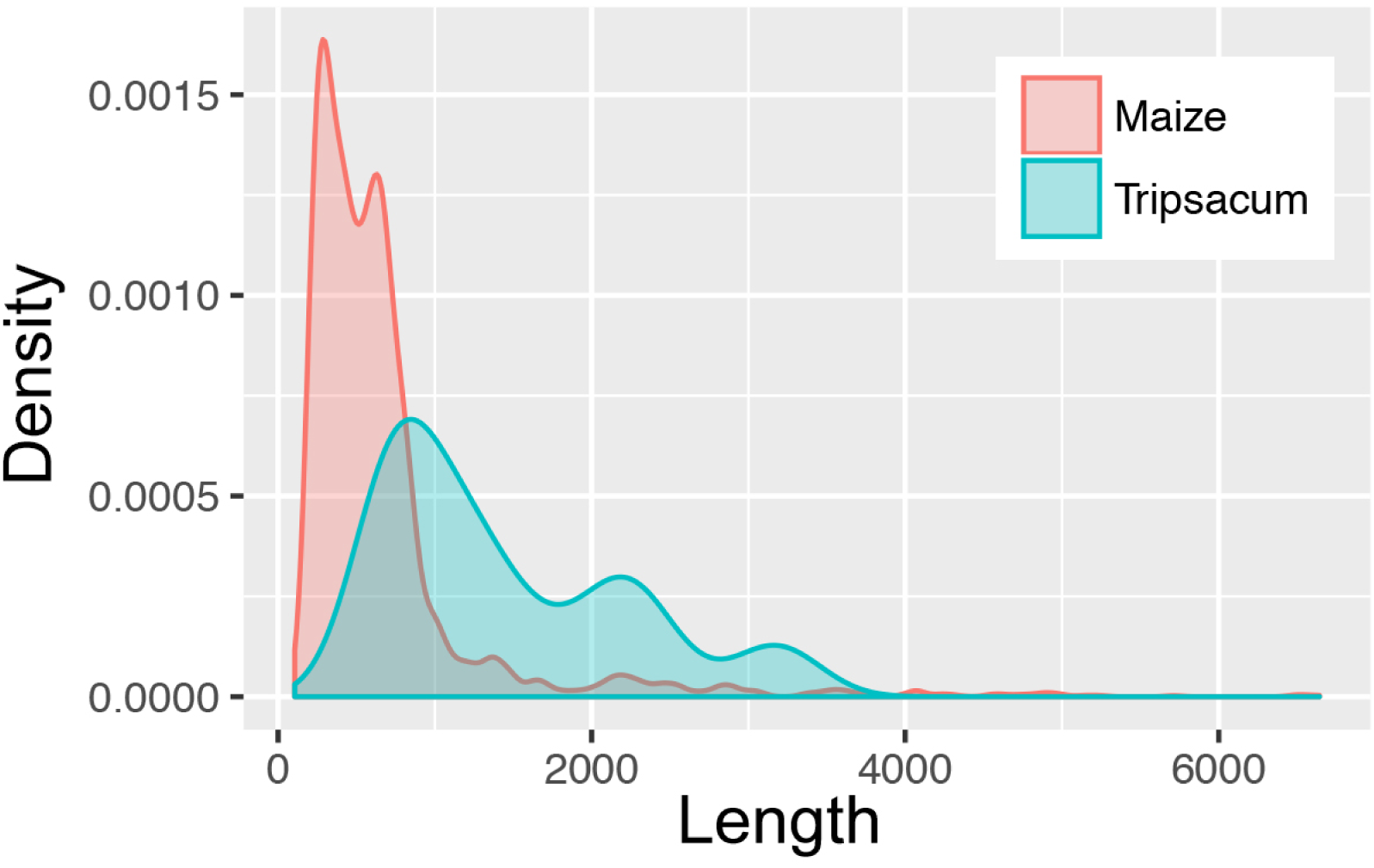
Comparison of length distribution of lncRNAs identified using Pacbio sequencing data between maize and tripsacum.

**Figure S8.**
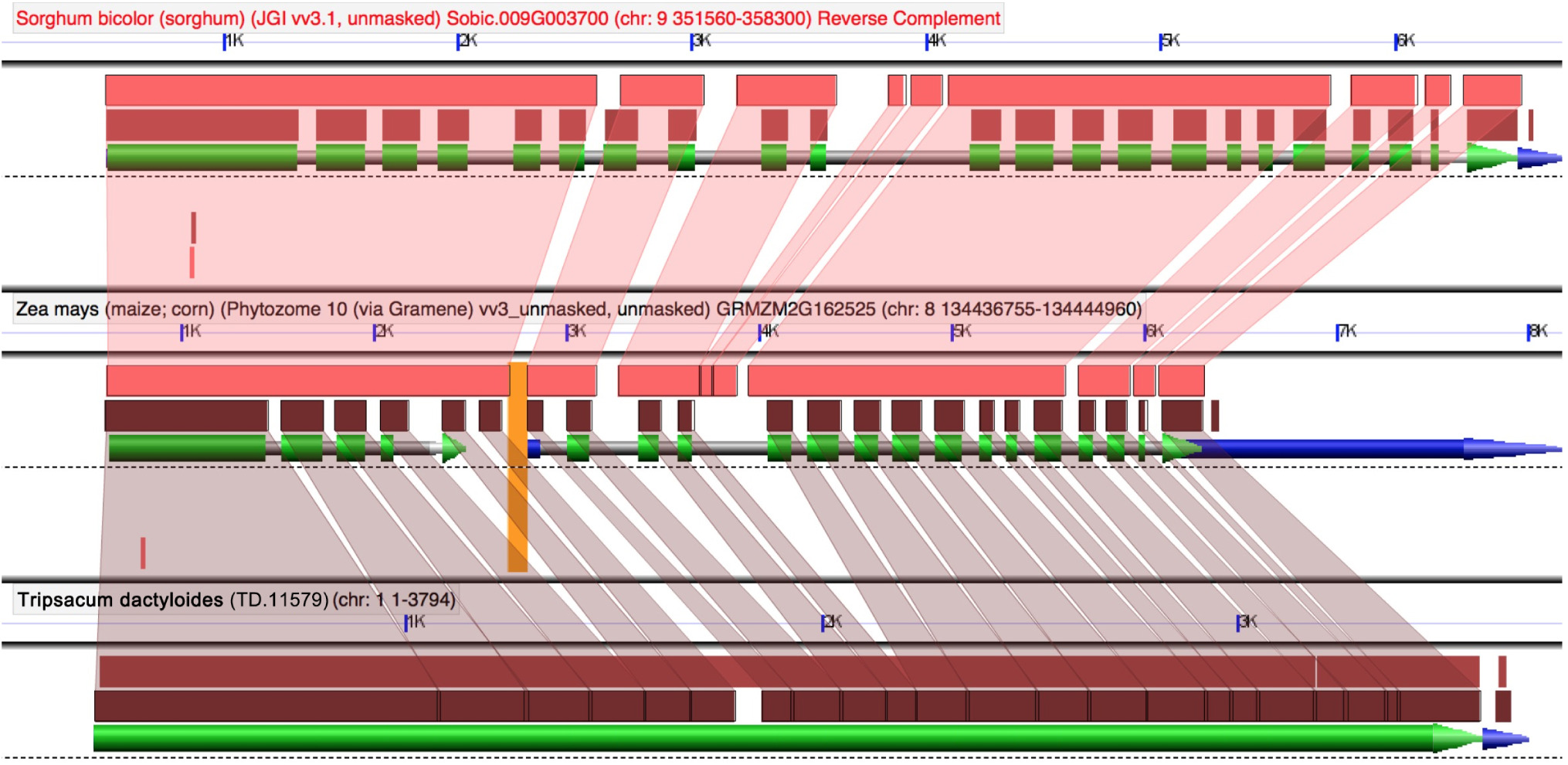
Example of one tripsacum transcript spans two maize gene models but correlated with one sorghum gene model through GEvo analysis.

**Figure S9.**
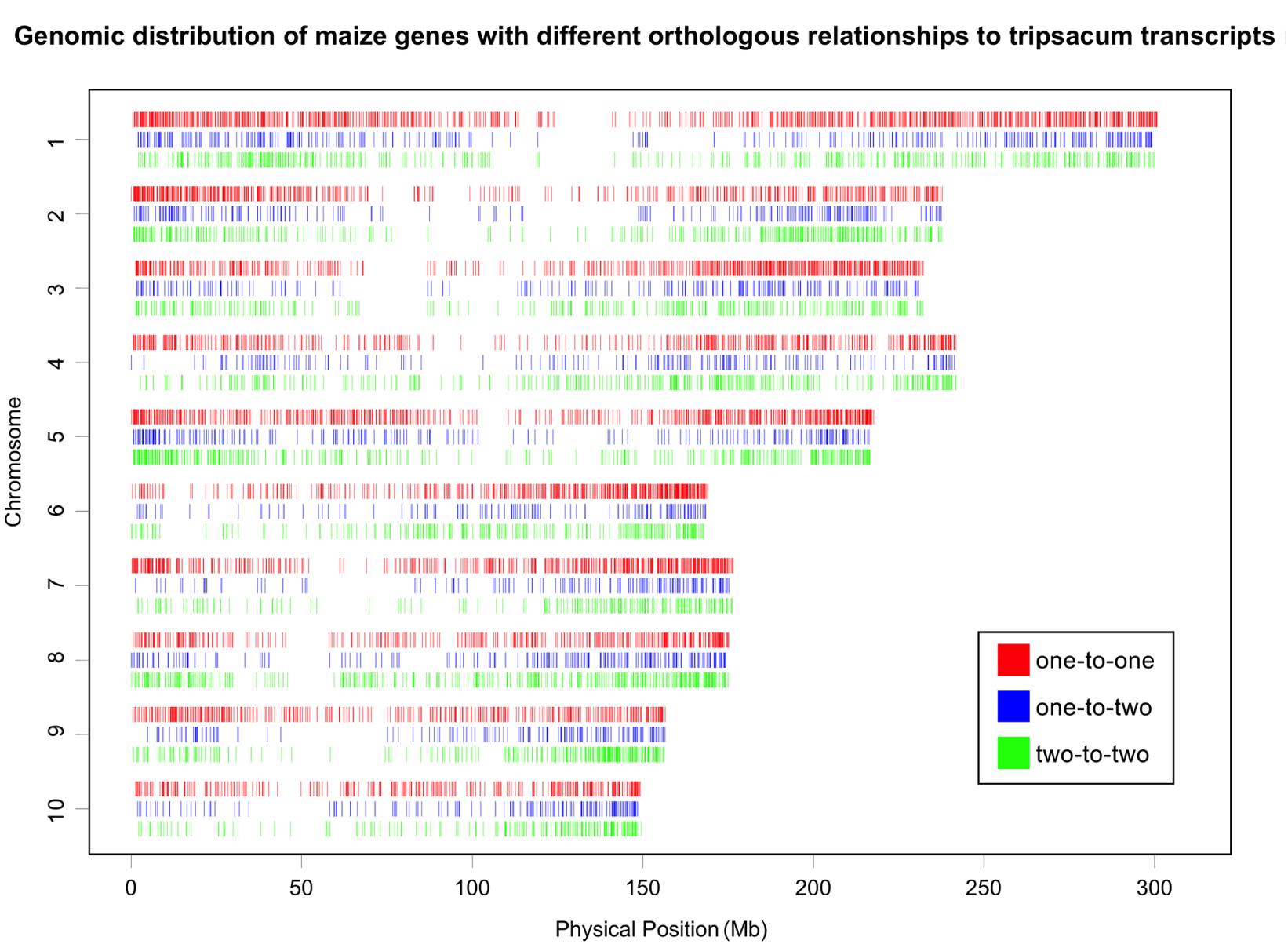
Distribution of maize genes with either 1:1, 2:1 or 2:2 relationships to assembled tripsacum transcripts across the maize genome.

**Figure S10.**
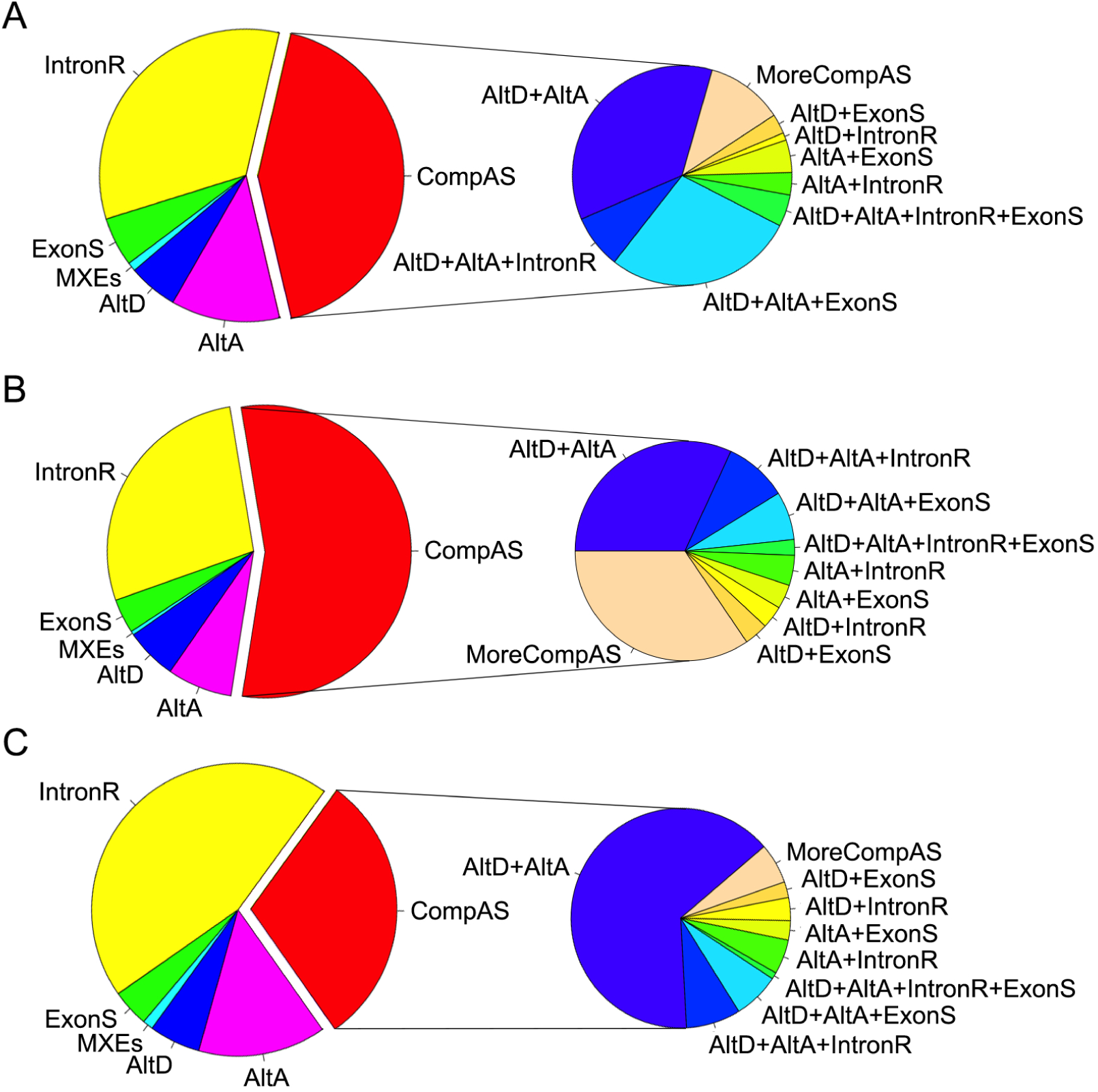
Proportions of alternative splicing (AS) types (Intron retention, IntronR; Exon skipping, ExonS; Alternative donor, AltD; Alternative acceptor, AltA; Mutually exclusive exons, MXEs; Complicated AS, CompAS; More complicated AS, MoreCompAS) found in (A) tripsacum and (B) maize using PacBio long sequences. (C) Proportion of conserved AS types between maize and tripsacum.

**Figure S11.**
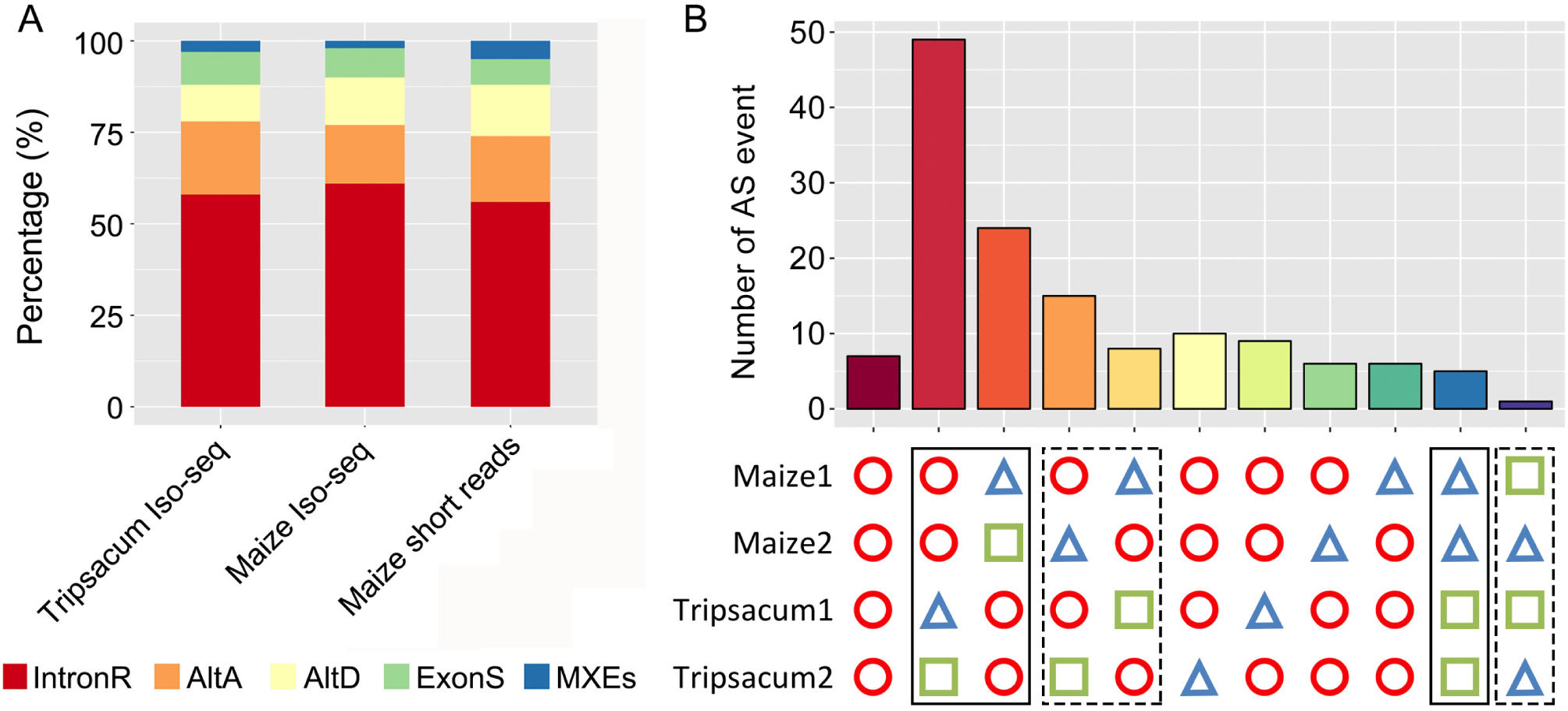
Comparison of alternative splicing (AS) distribution in (A) different species using long- and short-reads (long reads data for maize taken from [56] and short reads data of maize and sorghum were from [67]) and (B) subgenomes of maize and tripsacum. Identical shapes indicate genes with a conserved AS vent. Solid line boxes mark cases most parsimoniously explained by a change in *trans*-regulation of AS, while dashed line boxes mark cases most parsimoniously explained by a change in *cis*-regulation of AS.

